# A structure-based deep learning framework for protein engineering

**DOI:** 10.1101/833905

**Authors:** Raghav Shroff, Austin W. Cole, Barrett R. Morrow, Daniel J. Diaz, Isaac Donnell, Jimmy Gollihar, Andrew D. Ellington, Ross Thyer

## Abstract

While deep learning methods exist to guide protein optimization, examples of novel proteins generated with these techniques require *a priori* mutational data. Here we report a 3D convolutional neural network that associates amino acids with neighboring chemical microenvironments at state-of-the-art accuracy. This algorithm enables identification of novel gain-of-function mutations, and subsequent experiments confirm substantive phenotypic improvements in stability-associated phenotypes *in vivo* across three diverse proteins.

## Introduction

Protein engineering is a transformative approach in biotechnology and biomedicine commonly used to alter natural proteins to tolerate non-native environments^1^, modify substrate specificity^2^, and improve catalytic activity^3^. Underpinning these properties is a protein’s ability to fold and adopt a stable active configuration. This property is currently engineered either from sequence^4^, or energetic simulations^5^. Deep learning approaches have been reported, however these models either predict empirically measured stability effects in biased datasets containing only thousands of annotated observations^6^ or require model training on the target protein^7, 8^. Recently, a 3D-CNN was trained to associate local protein microenvironments with their central amino acid^9^. Given structural data, this model was able to predict wild type amino acids at positions where destabilizing mutations had been experimentally introduced. We hypothesized that the converse might also be true: stabilizing, gain-of-function mutations could be introduced at positions where the wild-type residue is disfavored. Here, we use a deep learning algorithm to improve *in vivo* protein functionality several fold by introducing mutations to better align proteins with amino acid-structure relationships gleaned from the entirety of the observed proteome.

## Results

In order to generate an algorithm that could identify unfavorable amino acid residues in virtually any protein structure, we trained a model to learn the correct association between an amino acid and its surrounding chemical environment, relying on the wealth of structures in the Protein Data Bank. We began by rebuilding the neural network architecture published by Torng and Altman with minor modifications (**Fig. 1a**, see **Online Methods** for details), replicating the reported classification accuracy of 41.2% (**Fig. 1b)** using the original training and testing sets (32,760 and 1601 structures, respectively)^9^. To improve the model’s performance, we made several discrete changes towards more explicit biophysical annotations adding in new atomic channels for hydrogen atoms and accommodating the partial charge and solvent accessibility for each atom, increasing accuracy to 43.4% and 52.4% respectively.

**Figure 1:**
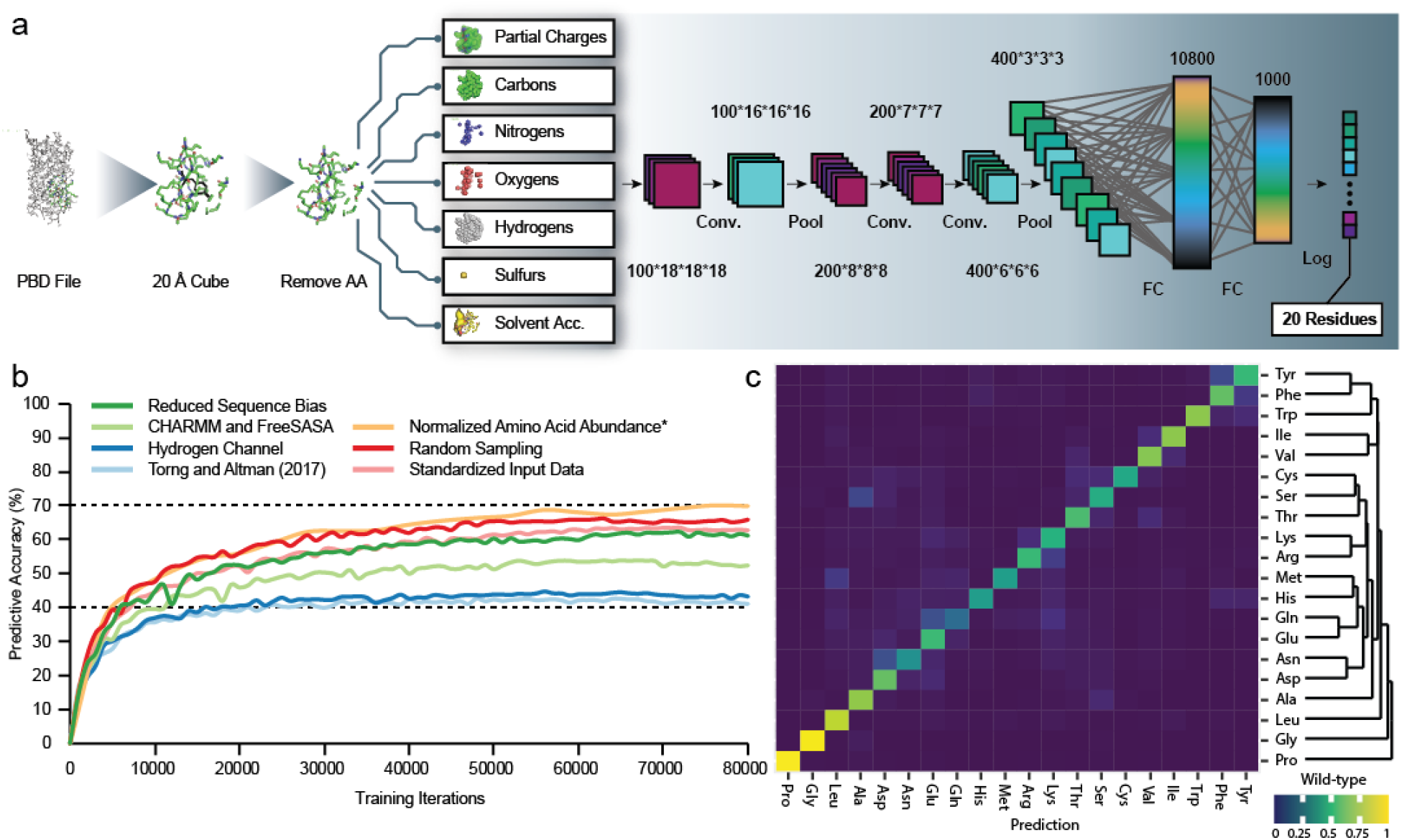
Design and performance of a deep learning program capable of classifying wild-type amino acids with improved accuracy. a) Schematic of the model depicting the data pipeline and neural network architecture. b) Discrete changes made to the neural net framework described by Torng and Altman^9^ and their effect on classification accuracy. *Normalizing the amino acid abundance of the training data increased the size of the dataset by roughly 4-fold. While the number of epochs decreased, the number of training iterations needed for convergence remained similar to the other versions. c) Confusion matrix showing bias of wild-type amino acid classification. Structurally unique amino acids Gly and Pro are assigned as wild-type with very high probability.

The selection methodology for both protein structures and amino acid residues introduced several biases to the training data. The dataset contained multiple structures of closely related proteins which biased training towards overrepresented protein structures, where the 32,760 PDB IDs map to only 11,418 UniProtKB IDs. Additionally, deposited crystallographic structures are refined by algorithms of their time which are not necessarily the current state of the art. To improve dataset composition and uniformity, we gathered all PDB structures with less than 2.5 angstrom resolution and at most 50% sequence similarity and drew from structures in the PDB-REDO database, where existing protein structures are refined in a uniform manner^10^. These two changes in data consistency resulted in 19,436 structures for training with an additional 300 structures for out-of-sample testing and increased wild-type prediction accuracy to 63%. Finally, by sampling amino acids randomly throughout a protein sequence and by mirroring the relative abundance amino acids in proteins represented in the PDB database (**SI Fig. 1**) the new training set, consisting of 1.6 million amino acid environments, improved classification accuracy to nearly 70%. In assessing a confusion matrix (**Fig. 1c**), similar amino acids were commonly misclassified, indicating that the neural network recapitulates known biochemistry and the genetic code. Furthermore, proline and glycine, which have unique structures, are classified with above 96% accuracy while glutamine is classified at only 33% accuracy.

We next investigated the performance of the model using empirical data from deep mutational scanning (DMS) experiments for the proteins TEM-1 β-lactamase, immunoglobulin binding domain of protein G (gb1), Aminoglycoside-3′-Phosphotransferase-Iia, ubiquitin, and Hsp90^11^. In this aggregate dataset, the effects of all possible single substitutions were quantified with a ceiling for activity set at wild-type function, i.e. no beneficial mutations were observable. We identified 292 positions where any substitution incurred a measurable fitness cost and benchmarked classification accuracy on this subset, the presumption being that the model’s classification accuracy on amino acids that are demonstrably best suited for a defined environment should exceed its overall classification accuracy. Consistent with this, the final version of the model achieves a recall of 87.0%, which is 25.4% higher than the starting model (**SI Fig. 2**), and 17% higher than its baseline out-of-sample accuracy. Precision recall curves for this task also confirm improvement over the starting point (**SI Fig. 3)**. Taken together, these data confirm our model classifies wild-type amino acids with unprecedented accuracy compared to previously reported deep learning approaches^9, 12–14^.

Having shown that we could accurately assign wild-type residues which are an optimal fit for their surroundings, we attempted to identify a series of gain-of-function mutations at wild-type amino acids assigned a low probability to occur in their native microenvironment. These disfavored wild-type residues might be substituted to improve fit within the panorama of natural protein structures and similarly improve folding and function of a protein. To assess phenotypes associated with folding and stability we investigated three model proteins: BFP, phosphomannose isomerase, and TEM-1 β-lactamase.

We initially tested this hypothesis in an engineered blue fluorescent protein secBFP2.1^15^ (**SI Table 1**) by building saturating libraries at residues assigned either the lowest (disfavored) or highest (favored) wild-type probabilities by our model. We also mutagenized ten residues selected at random to establish a control. Six of nine disfavored residues, one of ten random residues, and zero of ten favored residues could be substituted to improve fluorescence of secBFP2.1 (p = 0.01 by a Fisher’s exact test for disfavored versus random subsets; **Fig. 2a** and **SI Figs. 4-6**). We amalgamated the beneficial substitutions into a single variant, designated BFP-Bluebonnet (BB), which improved florescence in *E. coli* by more than six-fold (**Fig. 2b-c**). Furthermore, purified BFP-Bluebonnet exhibited improved thermal tolerance and chemical stability in guanidinium as compared to both secBFP2.1 and mTagBFP2 (**SI Fig. 7**). Blue fluorescent proteins are used less frequently than their counterparts across the visual spectrum for localization studies, e.g. GFP and RFP, in large part due to their maturation kinetics and solubility *in vivo*. Bluebonnet addresses these limitations and offers improved *in vivo* performance while retaining the advantageous secretory properties of its ancestor.

**Figure 2:**
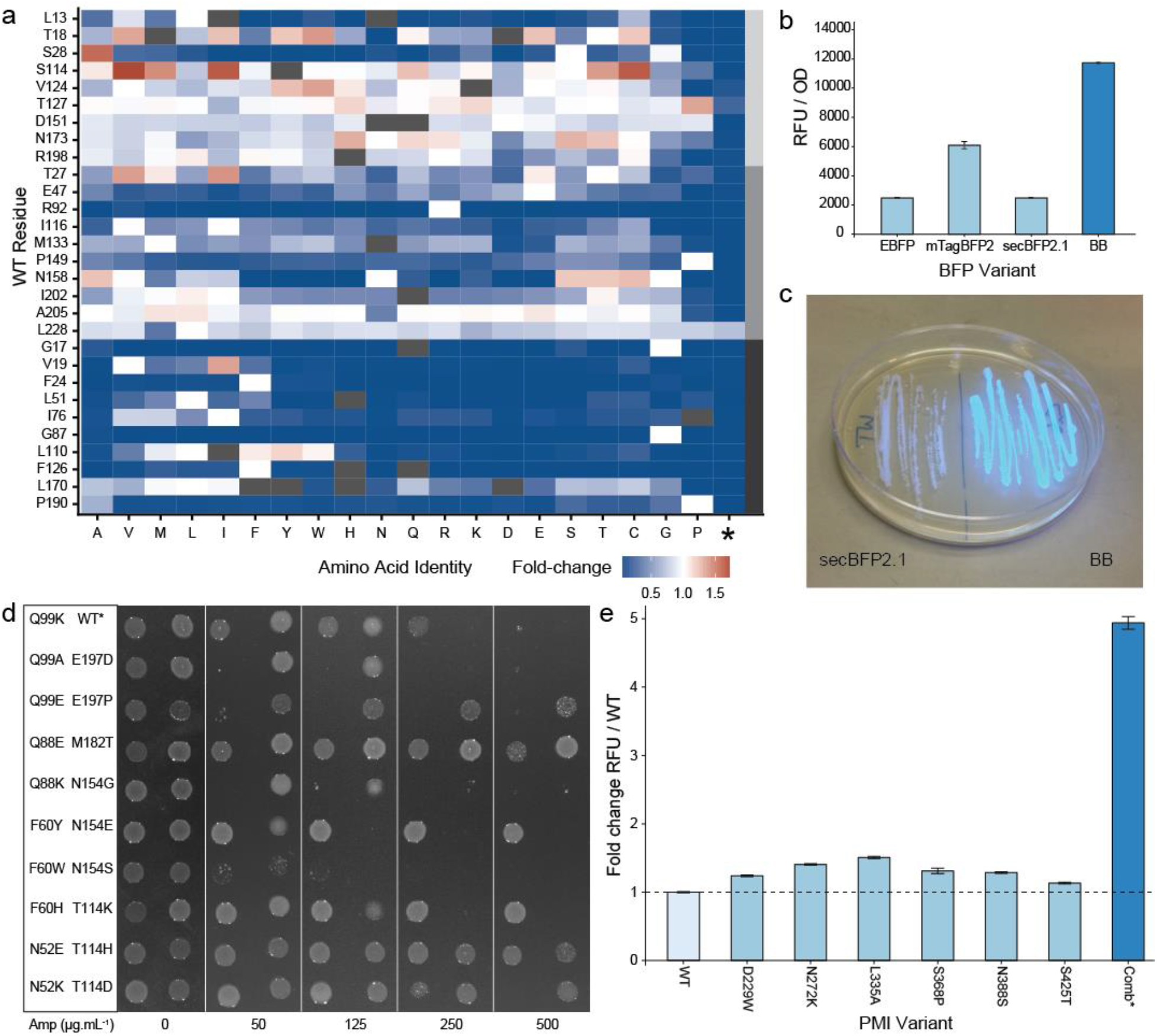
Empirical validation of the model as a tool for protein engineering using three model proteins. a) Heatmap showing fold-change over wild-type for site-saturation mutants of secBFP2.1. The light grey, dark grey and black bars on the right indicate the series of disfavored, random and favored residues respectively. Note, L228 is only five residues away from the C-terminus. Substitutions at this position, including stop codons, have minimal impact on fluorescence. b) An improved variant of secBFP2.1 containing mutations T18W, S28A, S114V, V124T, T127P, D151G, N173T and R198L. was ~6-fold more fluorescent in vivo than the parental protein. This variant was named BFP-Bluebonnet (BB). c) Plate assay showing increased in vivo fluorescence of BFP-Bluebonnet compared to secBFP2.1. d) Stabilizing mutations were identified in TEM-1 β-lactamase at N52, F60, Q88, Q99, T114, M182 and E197. WT* contains the destabilizing mutation L250Q. Residue Q88 was ranked as the 11^th^ least favorable in TEM-1 β-lactamase and was included in place of D214 which lies in the active site. e) Beneficial mutations were identified in CaPMI at residues D229, N272, L335, S368, N388 and S425. A combined mutant containing D229W, N272K, L335A, N388S and S425T was five-fold more fluorescent than wild-type using the split-GFP assay. While S368P was identified as stabilizing by itself, it was deleterious in combination.

To verify that our model was generalizable to catalytically active proteins, we built site-saturation libraries at the ten most disfavored residues in each of two structurally and functionally unrelated enzymes, TEM-1 β-lactamase and *Candida albicans* phosphomannose isomerase (*Ca*PMI). TEM-1 β-lactamase is a model protein for deep mutational scanning and protein evolution, thereby providing a rich benchmark of cross-validated mutational annotations for the 3D CNN’s predictions. *Ca*PMI is poorly soluble in *E. coli* and lacks an easily screened readout for directed evolution, and thereby serves as an exemplar of how the model can be applied in conjunction with a generalizable high-throughput protein engineering workflow (see **Online Methods**). Seven of the ten residues in TEM-1 β-lactamase and six of the ten residues in *Ca*PMI could be substituted to improve phenotypes associated with folding and stability (**Fig. 2d-e**). Aggregating these stabilizing mutations improved the folding of *Ca*PMI in *E. coli* by five-fold without abolishing catalytic activity (**SI Fig. 8**).

To further simulate a situation in which large-scale screens are not possible, we examined the ability of the model to directly predict the beneficial substitutions for positions where the wild-type residue was assigned a low probability. Using this approach, more than 20% of directly predicted mutations improved the functional readout and furthermore, mutational effects additively improved each of the three distinct phenotypes at least five-fold in combination (see **Supporting Results** and **SI Figs. 9-11**). Additionally, the assignment of probabilities for all amino acids at a given position provides opportunities to explore fundamental aspects of protein structure and function. Using our model, we interrogated the microenvironment surrounding Met 182 in TEM-1 β-lactamase where a Met to Thr substitution results in global stabilization. We identified key backbone atoms which favor the Thr substitution in agreement with the findings of previous MD work at this locus^16^ (see **Supporting Results** and **SI Fig. 12**).

Two well-documented, alternative computational approaches to guide protein stabilization are Rosetta pmut_scan and FoldX PositionScan, both of which rely on energetics simulations. If our model learned inferences accessible by energetics calculations in either of these programs, we would expect significant overlap between the disfavored residues it identified and destabilizing positions predicted by either of these programs. Only three of thirty positions identified by the model were also identified by either Rosetta or FoldX, which also largely identified separate residues. Furthermore, in TEM-1 β-lactamase, each of these methods uniquely identified stabilizing mutations reported elsewhere in the literature^17, 18^ (**SI Fig. 13**). Therefore, our model can identify novel stabilizing loci not captured by other commonly used programs.

## Discussion

Here we report a modified 3D CNN architecture with state-of-the-art classification accuracy for assigning wild-type residues throughout proteins. Where native amino acids deviate from their structural and chemical consensus, we demonstrate that these positions with low wild-type favorability are excellent targets for site-saturation mutagenesis and yield stabilizing mutants at frequencies that exceed random selection. Combining the stabilizing mutations identified in three model proteins improved variant phenotypes several fold relative to their ancestor. Furthermore, this model is synergistic with existing protein design tools by identifying sets of mutations that do not overlap with those derived from energetics simulations. This work is the first demonstration of using deep learning to empirically improve protein function and opens a new avenue for protein engineering.

This tool is freely available for academic use at www.mutcompute.com.

### Methods

Detailed methods are available in the **Online Methods**.

## Acknowledgements

Funding from the Welch Foundation (F-1654), the ARO (SP0036191-PROJ0009952 via MURI), and the NSF (1541244) is acknowledged.

## Author Contributions

RS, AC, JG, AE and RT designed the experiments. RS, AC and DD wrote the code for the deep learning algorithm. RS, AC, BM, ID and RT performed the experimental work. RS, AC, AE and RT wrote the manuscript.

## Online Methods: A structure-based deep learning framework for protein engineering

### Computational Methods

#### Dataset

To reduce any bias resulting from the differential abundance of protein families in the PDB, we sought to build a dataset of protein structures with balanced phylogeny. To achieve this, we took all structures in the PDB database and clustered to 50% similarity to avoid oversampling towards certain protein classes. We further reduced the variability in the dataset by cross-referencing the structures to the PDB-redo database^1^, which uses a consistent algorithm to refine, rebuild, and validate structures from raw crystallographic data. Within each clustered set of sequences, we identified the structure with the lowest resolution. If no structure existed below a resolution of 2.5 angstroms the entire cluster was discarded. This process yielded 19436 structures, of which 300 were randomly set aside for out of sample testing and the remainder used to generate the training set.

#### Box Extraction

In addition to atomic annotations, our model adds additional channels for the partial charges and solvent accessibility associated with each atom. While all structure files label oxygen, carbon, nitrogen, and sulfur, hydrogens may be missing depending on the resolution of the structure. Using the program pdb2pqr (v2.2.1)^2^, hydrogens were placed into the structure and optimized while partial charges were assigned with the CHARMM force field. Solvent accessibility was calculated with the program FreeSASA (v2.0.2)^3^. To avoid oversampling residues from larger proteins, we limited the number of sampled environments from an individual protein to either half of the length of the protein or 100 amino acids, whichever number was less. Atomic environments consisting of a 20 angstrom cube centered around a single residue were generated as described in Torng and Altman^4^.

#### Neural Network Training

The convolutional neural network was built using theano (v1.0.3) and consists of six layers, all with ReLu (rectified linear unit) activations. The first two convolutions were performed with a filter size of 3×3×3 with no padding and increased the depth to 200 channels. We then performed a max pooling step, followed by two additional convolutions with a filter size of 2×2×2 and increasing the depth to 400. Max pooling was used again before flattening and feeding into two successive fully connected layers with dropout rates of 0.5 and 0.2, respectively. Softmax activation was applied to the logits to obtain probability scores for each of the 20 amino acids.

Neural network training was performed on TACC’s Maverick cluster with a NVIDIA Tesla K40 GPU. 1.6 million amino acid environments were generated with the abundance of individual amino acids mirroring the natural frequency observed in the PDB. As the dataset was too large to load entirely into memory, we split the data into 20,000 samples and randomly shuffled the order after loading. Batch sizes of 20 samples were used and the loss was calculated through RMSprop. Training was performed with an adaptive learning rate and lowered by 10% if validation accuracy did not decrease within 8000 training iterations. Four epochs were run, at which point overfitting was observed. Test and validation accuracy were measured in 6000 amino acid environments with equal representation of each residue.

#### Confusion Matrix and Regression Bias

To calculate the frequency at which wild-type residues were correctly predicted, 20,000 amino acid environments were generated from out of sample PDBs (i.e. structures not seen during training) with an amino acid distribution mirroring natural frequencies. Regressions highlighting amino acid bias were created by plotting the sum of the predicted probability values against the frequency in the test set. The confusion matrix was generated by plotting the single amino acid assigned the highest probability at each microenvironment sampled compared to the wild-type amino acid.

#### Rosetta/FoldX Calculations

The pmut_scan program within the Rosetta software suite (v3.9) was used to calculate the computational effect of mutations with a large ΔΔG cutoff value to output both stabilizing and destabilizing mutations. To perform the analogous operation in FoldX (ver. 4), the PositionScan module was used. In either program, the least favorable sites were found by summing values less than zero (the sign change of a stabilizing mutation) and identifying the ten sites with the most negative value.

#### Deep Mutational Scanning Analysis

Each computational method was assessed using deep mutational scanning data sets paired with the corresponding structures: TEM-1 β-lactamase, PDB:1BTL; protein G, PBD:2QMT; aminoglycoside-3’-phosphotransferase-IIa, PDB:1ND4; ubiqutin, PDB:4XOF, and Hsp90, PDB:2BRC. Normalized fitness values were derived from Gray *et al*. (2018)^5^ with a threshold of 1.02 to determine if a variant greater than wild-type exists. Within this subset, a positive result was defined if no other variant empirically exhibited better fitness than wild-type and, for our model, the wild-type amino acid was assigned the largest probability, or, for Rosetta and FoldX calculations, the minimum ΔΔG value (i.e. the most stabilizing value) was greater than zero.

### Molecular Methods

#### Molecular Biology

Experiments described in this manuscript were performed using standard molecular biology techniques. Unless otherwise indicated, all plasmids, single point mutations in reporter genes and site-saturation mutagenesis libraries were constructed using Gibson assembly. For site-saturation mutagenesis libraries, 2 μL of the reaction mixture was transformed into 50 μL of chemically competent *E. coli* cells. Transformations were required to exceed 10-fold library coverage (> 320 single colonies).

#### BFP Fluorescence Assay

SecBFP2.1 was cloned into a kanamycin resistant derivative of plasmid pQE flanked by a T7 promoter and terminator. Site-saturation libraries were transformed into *E. coli* strain BL21 DE3 and a series of 10-fold dilutions (spanning two orders of magnitude) were plated on solid media to ensure sufficient discrete single colonies. 96-well deep-well plates were inoculated with 92 individual library transformants and four wild-type controls. Two plates were assayed for each library. Cells were cultured ON at 37 °C in plate shakers at 850 rpm. 20 μL of the ON cultures were diluted into 880 μL LB and incubated for two hours. Cells were induced by the addition of 100 μL media containing 0.5 mM IPTG, resulting in a final concentration of 50 μM. After a four hour induction, cells were harvested by centrifugation and resuspended in 1 mL PBS. Fluorescence was measured on a Tecan M200 Pro using 400 nm for excitation and 460 nm for emission. A maximum of 12 individuals at each library site exhibiting fluorescence / OD600 values greater than wild-type were sequenced. Candidate mutations were re-cloned into the pQE plasmid and rephenotyped. Rephenotyping was performed in biological and technical triplicate.

#### Protein Purification

To purify secBFP2.1 mutants, a 6xHis tag was appended to the C-terminus via a Gly-Ser-Gly linker. BL21 DE3 cells were cultured in Superior Broth to mid-log phase (~ OD600 0.6) and induced with 1 mM IPTG for 16 hours at 18 °C. Following induction, cells were harvested by centrifugation and lysed by sonication in 50 mM sodium phosphate, 300mM NaCl, 20mM Imidazole pH 7.4 buffer containing protease inhibitor (Pierce Protease Inhibitor) and Benzonase Nuclease (EMD Millipore). Cell lysate was clarified by centrifugation (40000 x *g*) and BFP variants purified using HisPur™ Ni-NTA Resin. Purified protein was dialyzed into 50 mM sodium phosphate pH 7.4 buffer and analyzed by SDS PAGE to assess purity.

#### Thermal Melt Assay

Purified blue fluorescent proteins were diluted to 0.01 mg.mL^-1^ in PBS pH 7.4 and 100 uL aliquots were heat treated for 10 minutes in PCR strips on a thermal gradient using a thermal cycler. Fluorescence of thermally challenged variants and controls incubated at room temperature was assayed using excitation and emission wavelengths of 402 nm and 457 nm respectively. Fluorescence readings were normalized to the mean of solutions incubated at room temperature e.g. a measurement of 0.8 indicates that a heat treated protein retained 80% of its untreated fluorescence.

#### Guanidinium Denaturation Assay

Purified blue fluorescent proteins were diluted to 0.01 mg.mL^-1^ in 6 M guanidinium hydrochloride. 100 uL aliquots in technical triplicate were added to wells of a 96-well clear-bottom black-walled plate and incubated at 25 °C for 23 hours. These purified fluorescent proteins were assayed at 30 minute intervals using excitation and emission wavelengths of 402 nm and 457 nm respectively. Plates were agitated preceding each measurement. Fluorescence values measured at time zero were used to normalize fluorescence through the remainder of the assay e.g. a measurement of 0.8 indicates that the protein retained 80% of its initial fluorescence.

#### TEM-1 Assay

The *bla*TEM-1 gene encoding TEM-1 β-lactamase, including the native promoter, was amplified from pETDuet-1 and cloned into pCDFDuet-1 immediately upstream of the second T7 terminator, replacing both T7 promoters and both polylinkers. The L250Q mutation was introduced into TEM-1 to destabilize the protein and enable easy identification of compensatory stabilizing mutations^6^. Site-saturation libraries were transformed into *E. coli* strain DH10B and recovered ON in liquid medium supplemented with spectinomycin. ON cultures were diluted and plated on a range of different carbenicillin concentrations (0, 50, 125, 250 and 500 μg.mL^-1^). For each library, 12 single colonies from the plate containing the highest concentration of carbenicillin were isolated and the *bla*TEM-1 gene sequenced. Beta-lactamase variants identified by library screening were recloned into pCDFDuet-1 and rephenotyped. Rephenotyping was performed by diluting overnight cultures, in biological triplicate, 100-fold and spotting 5 μL onto solid media containing a gradient of carbenicillin concentrations.

#### PMI Stability Assay

Improved variants of *Ca*PMI were identified using the split GFP reporter system described by Cabantous *et al*. (2008) with minor modifications^7^. Briefly, a fusion protein consisting of residues 173-238 of folding reporter GFP, a (GGGS)2 linker, residues 2-440 of *Ca*PMI, a (GGGS)2 linker and residues 2-172 of superfolder GFP was assembled in a derivative of pACYCDuet-1 (**SI Table 1**). Site-saturation libraries were transformed into *E. coli* strain BL21 DE3 and a series of dilutions plated on solid media supplemented with 0.25 mM IPTG. Following ON incubation at 37 °C, plates were further incubated at 4 °C for eight hours at which point highly fluorescent colonies were manually selected. PMI variants were subcloned and the fusion protein ORF fully sequenced prior to rephenotyping to ensure that increased fluorescence was not the result of mutations in the GFP fragments, linker regions, or plasmid backbone. Transformants were screened as described for BFP in 96-well deep-well plates in biological and technical triplicate. Fluorescence was measured on a Tecan M200 Pro using 475 nm for excitation and 535 nm for emission.

#### PMI Functional Assay

The *manA* gene encoding phosphomannose isomerase was disrupted in *E. coli* strain BL21 DE3 using lambda red recombineering to introduce a kanamycin resistance marker. Successful deletions were confirmed by colony PCR of Kan^R^ colonies using primers which flanked the *manA* locus. The wild-type *C. albicans manA* gene or variants containing combinations of stabilizing point mutations were cloned into a derivative of pACYCDuet-1 using Gibson assembly. BL21 DE3 Δ*manA*::kan cells were transformed with PMI expression plasmids and plated on LB agar with appropriate antibiotics. Single transformants, in biological triplicate, were transferred to liquid M9 minimal medium with 0.4% glucose and cultured ON. Cells were washed in a 1:1 volume of M9 medium without any carbon source and 2 μL streaked on M9 minimal medium plates supplemented with 0.4% mannose and 0.25 mM IPTG. Wild-type BL21 DE3 cells and BL21 DE3 Δ*manA*::kan cells containing an empty expression plasmid were used as positive and negative controls respectively. Plates were incubated at 37 °C for 24 hours.

#### NGS

Purified plasmids encoding BFP variants were quantified using the QuantIT dsDNA Assay Kit (Thermo Scientific) and 50 ng of each plasmid was pooled by well position, resulting in 192 samples each with a different variant for each tested position. Samples were prepped for sequencing by amplifying two discrete ~350 bp regions of the BFP gene with primers containing Illumina adapters and dual indexes. Sequencing was performed using Illumina MiSeq with paired end 2×300 bp reads.

#### Variant Calling

Following sequencing, paired end reads were joined together using fastq-join (v1.3.1). Raw sequences were aligned to the original gene sequence with bwa (v0.7.17). Read counts across a single position were normalized to the observed fraction of each codon. Variants that contained at least 100 read counts and exceeding 1% of wild-type counts were labeled as such, while variants that failed were left unlabeled. Outliers were identified through the OPTICS algorithm in the scikit-learn package (v0.21.3). Each sample was normalized by dividing the fluorescence by the OD600 and as well as to the average wild-type value per plate. Significance was determined using a Fisher’s exact test where success of a position was defined as the mean fluorescence of the most fluorescent variant being at least three standard deviations higher than the wild-type mean.

### Statistical Methods and Data Presentation

All data in the manuscript are displayed as mean ± s.e.m. unless specifically indicated. Bar graphs, regressions, confusion matrix, NGS variant graphs were plotted in R 3.4.1 using the package ggplot2 (v2.2.1).

## Supplementary Information: A structure-based deep learning framework for protein engineering

### Supplementary Results

While site-saturation mutagenesis at candidate residues is a good option for identifying beneficial mutations, it relies on the ability to screen protein variants in at least moderate throughput. To simulate a situation in which screening at such a scale is not possible, we examined the ability of the model to directly identify beneficial substitutions at residues where it did not assign the highest probability to the wild-type amino acid. We built all unique single point mutations in the three proteins (secBFP2.1, TEM-1 β-lactamase and *Ca*PMI) using the top ten substitutions generated by three different but largely overlapping interpretations of the model: the amino acid assigned the highest probability at the residues with lowest assigned probability of the wild-type amino acid (the residues selected for site-saturation mutagenesis), the amino acid assigned the highest probability where it differed from wild-type (regardless of the wild-type probability), and mutation to the amino acid assigned the highest probability differing from wild-type resulting in the greatest log-fold change over the wild-type probability (**SI Figs. 9-11**). Using this approach, several individual stabilizing mutations were identified for each protein and the effects were largely additive when combined. Although no single interpretation of the output data was clearly superior, this methodology resulted in at most 22 unique variants, which is a manageable number to synthesize and screen for all but the most challenging proteins.

Although we focused on protein engineering applications for our model, it also has considerable potential as a tool to unravel fundamental biology. In particular, we sought to explain the model’s ability to flag the mutation M182T in TEM-1 β-lactamase, a global suppressor mutation that has been identified in many clinical isolates. Despite its identification decades ago, the mechanistic explanation for stabilization remains under debate. One model proposes that the threonine hydroxyl forms an N-cap H bond with Ala 185 as determined through crystallographic analysis^1^, while a competing explanation determined through molecular modeling suggests a stabilizing hydrogen bond with Glu 63 and/or Glu 64^2–4^. To find the contributing atoms that most favor a mutation to threonine, we systematically deleted every atom in the Met 182 microenvironment and used the used the model to analyze where the probabilities changed the most. Our method flagged two atoms, the backbone oxygen of Glu 63 and the amide hydrogen on Ala 185, in which the removal of either atom decreased the probability of observing a threonine by over 200 fold (SI Figure 12). Thus, a neural network framework can be used to suggest stabilization mechanisms in addition to identifying candidate residues for mutagenesis.

### Supplementary Figures

**SI Figure 1:**
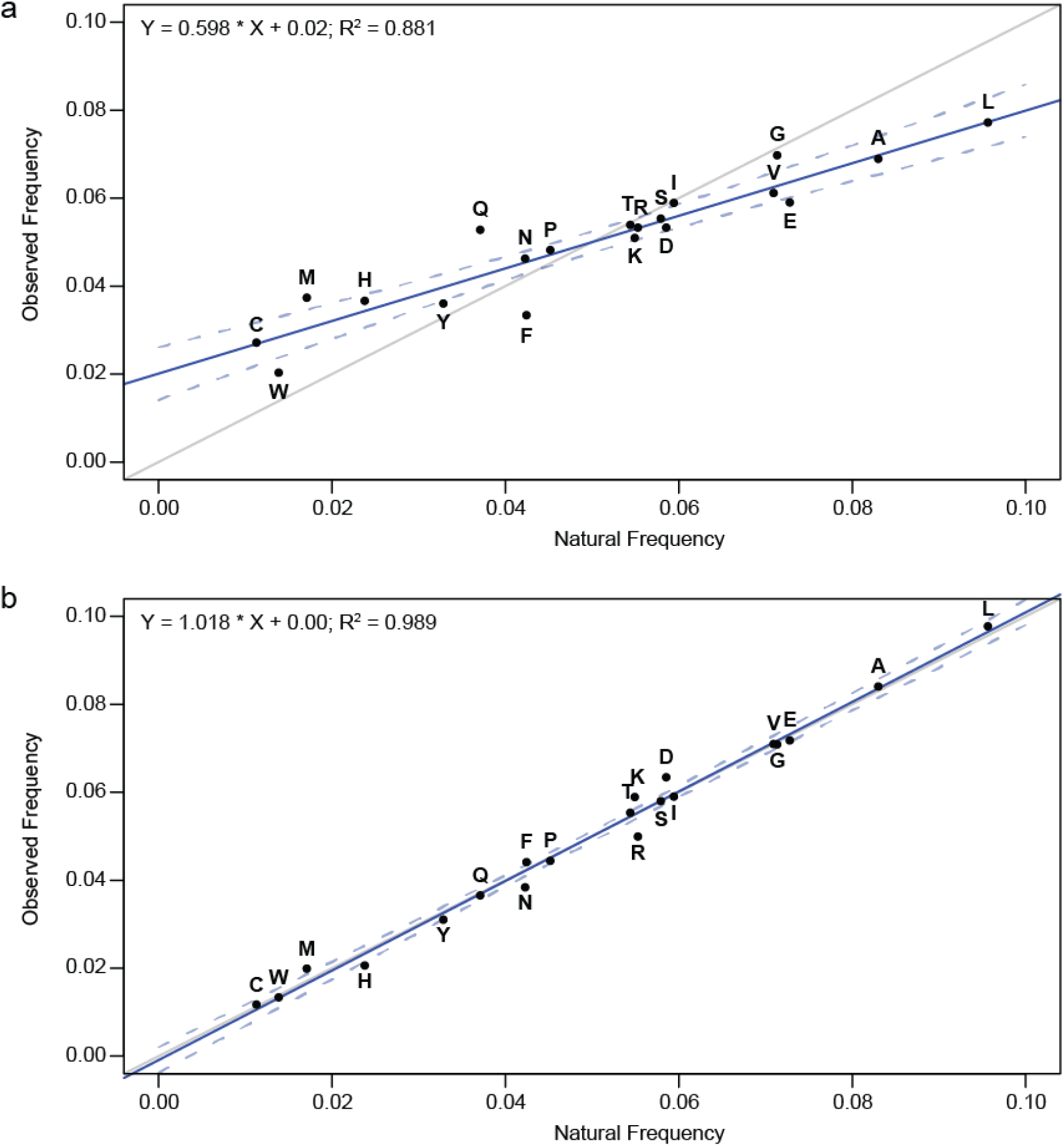
Improvement in predictive accuracy of the model when training was normalized for amino acid abundance. a) Classification accuracy of the model trained for 84,000 iterations without normalization. The p-value for a two-sided t-test against null hypothesis: ‘the slope is equal to 1’ is less than 10e-6. b) Classification accuracy of the model following training for 240,000 iterations correcting for amino acid abundance. The grey line depicts a line with slope 1 and the blue line is the regression for the observed amino frequencies compared to the natural abundance. Dotted lines delineate 95% confidence intervals for the regression. The p-value for a two-sided t-test against null hypothesis: ‘the slope is equal to 1’ is 0.48.

**Figure 2:**
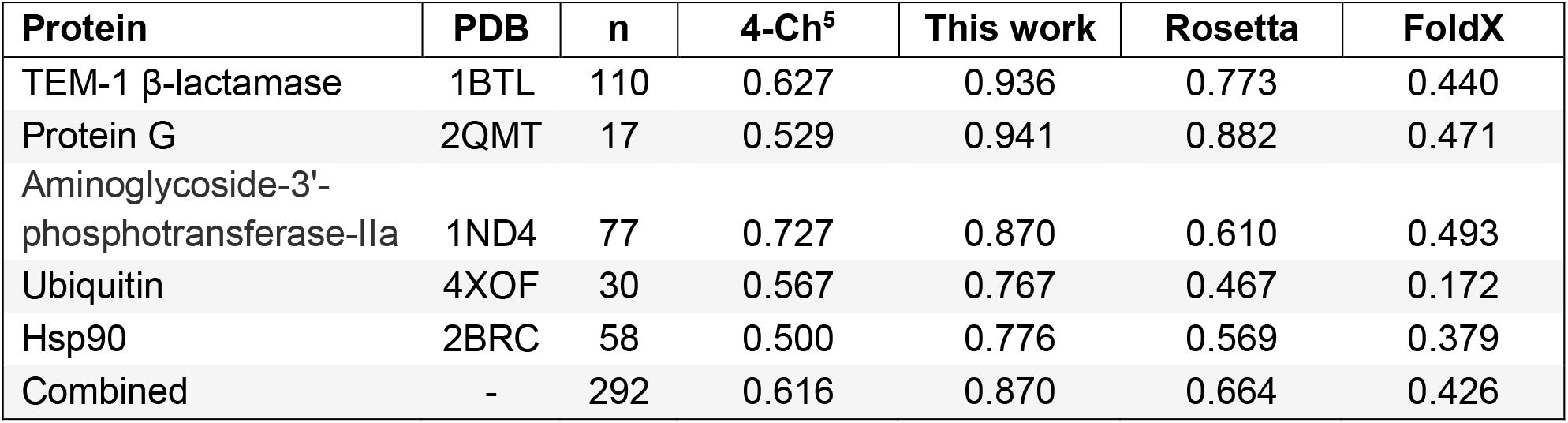
Classification accuracy of different computation tools for protein engineering using a dataset of true positive wild-type residues. For the 4-channel model and the model presented in this work, classification was considered correct if the wild-type amino acid was assigned the highest probability. For Rosetta and FoldX, classification was considered correct if the wild-type amino acid was assigned the lowest ΔΔG.

**SI Figure 3:**
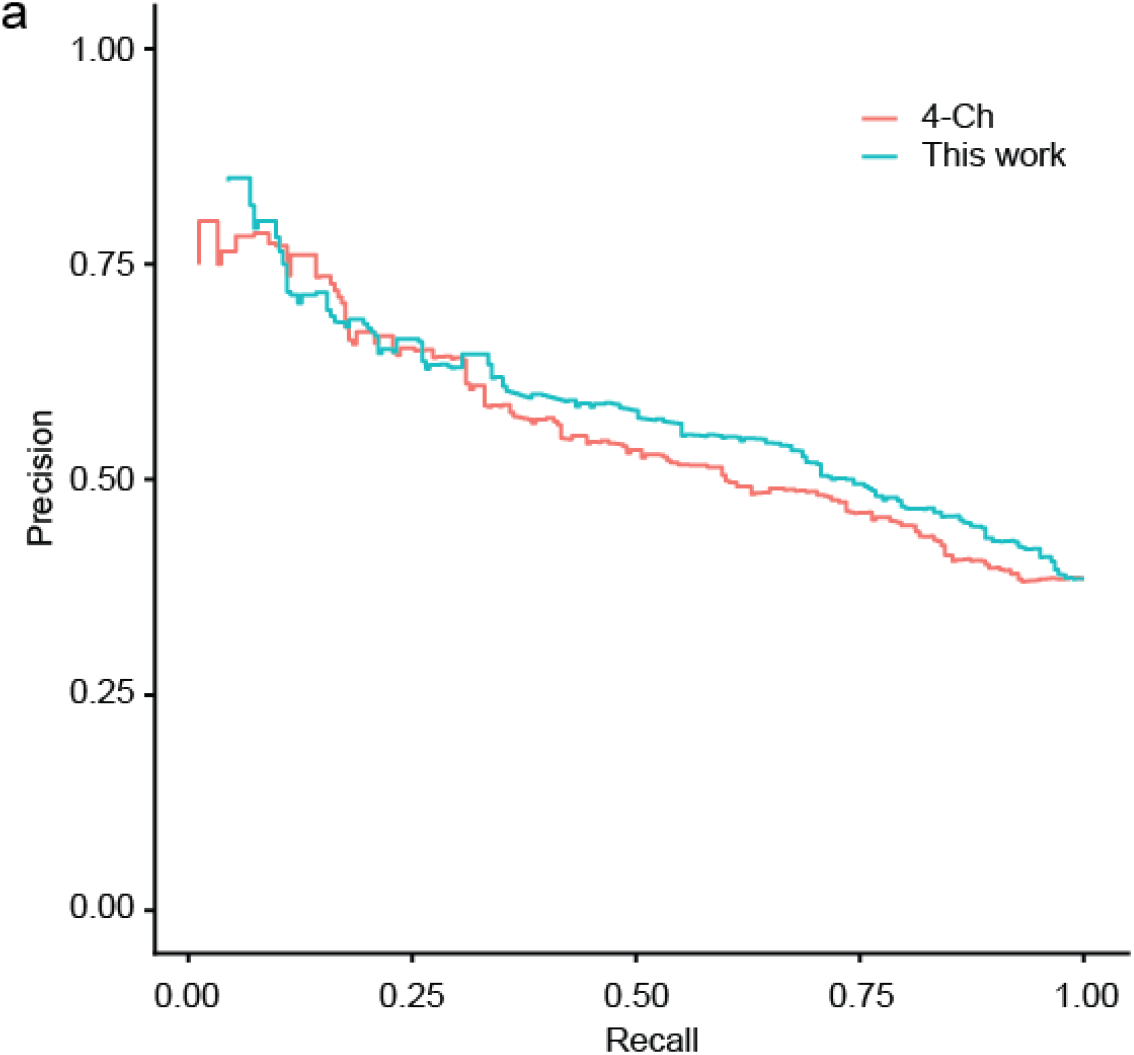
Precision recall curve of the initial and improved deep learning models. Positions where the wild-type residue exhibited the greatest normalized fitness were aggregated from five deep mutational scanning (DMS) datasets. Input data was drawn from DMS datasets and PBD files for the following proteins: TEM-1 β-lactamase, PDB:1BTL; protein G, PBD:2QMT; aminoglycoside-3-phosphotransferase-IIa, PDB:1ND4; ubiqutin, PDB:4XOF, and Hsp90, PDB:2BRC. The ability of both methods to identify these positions as wild-type was analyzed as the threshold for classification was varied.

**SI Figure 4:**
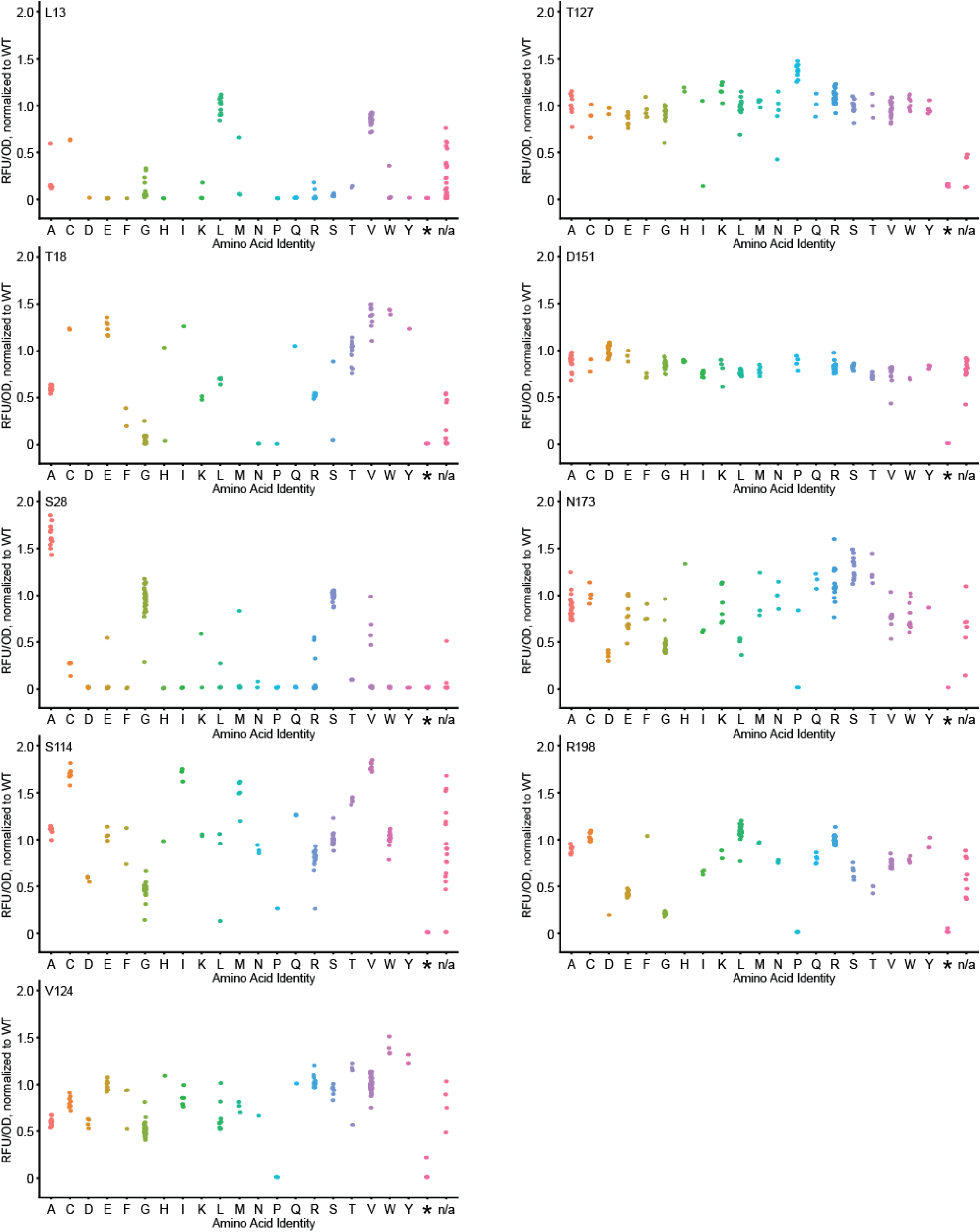
Fluorescence data for site-saturation libraries at disfavored residues in secBFP2.1 identified by the model. Raw fluorescence values were normalized to OD600 and to the average wild-type value. Outliers were identified through the OPTICS algorithm and removed. n/a represents variant calls that failed to meet the specified thresholds (see Methods).

**SI Figure 5:**
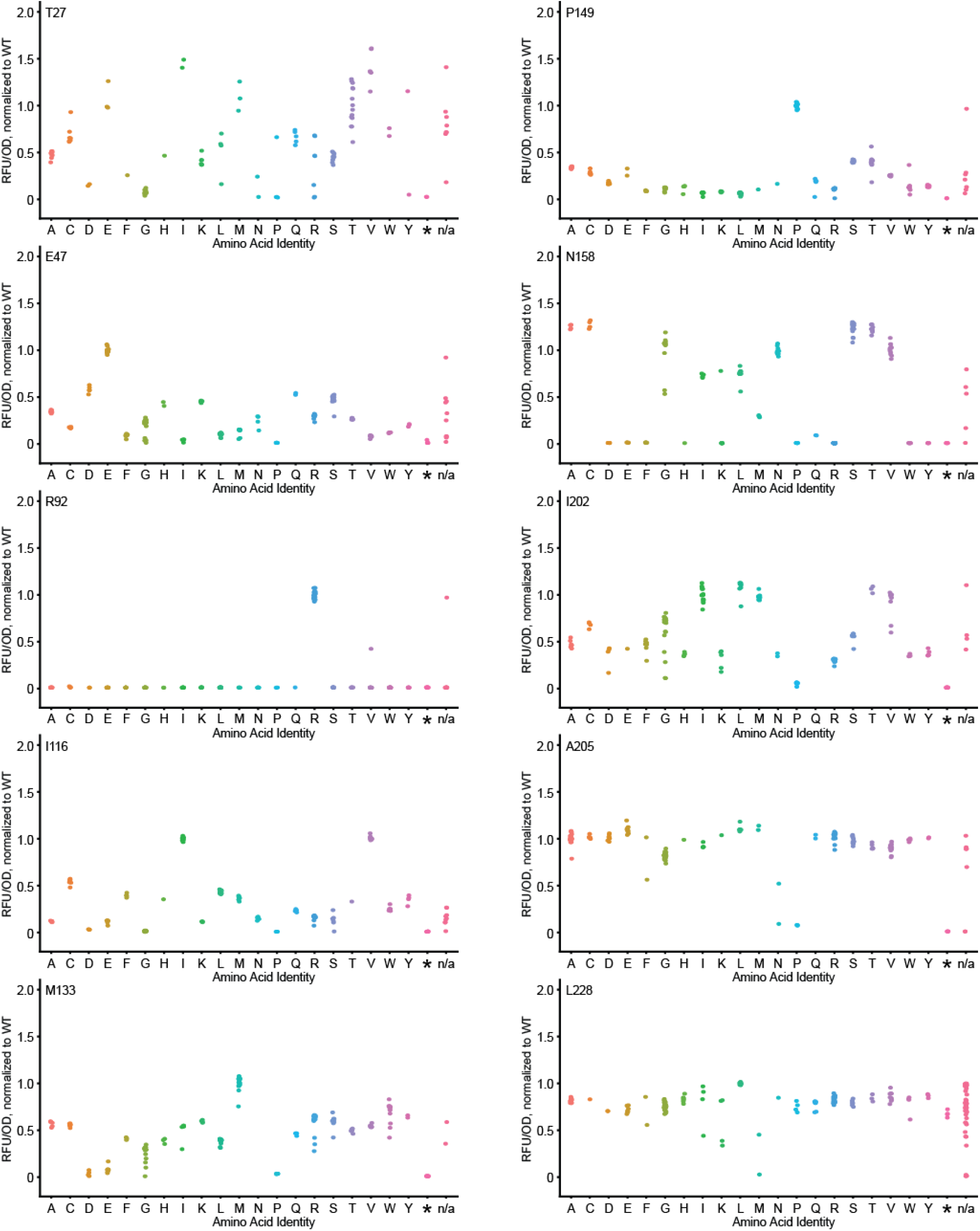
Fluorescence data for site-saturation libraries at random locations in secBFP2.1. Raw fluorescence values were normalized to OD600 and to the average wild-type value. Outliers were identified through the OPTICS algorithm and removed. n/a represents variant calls that failed to meet the specified thresholds (see Methods).

**SI Figure 6:**
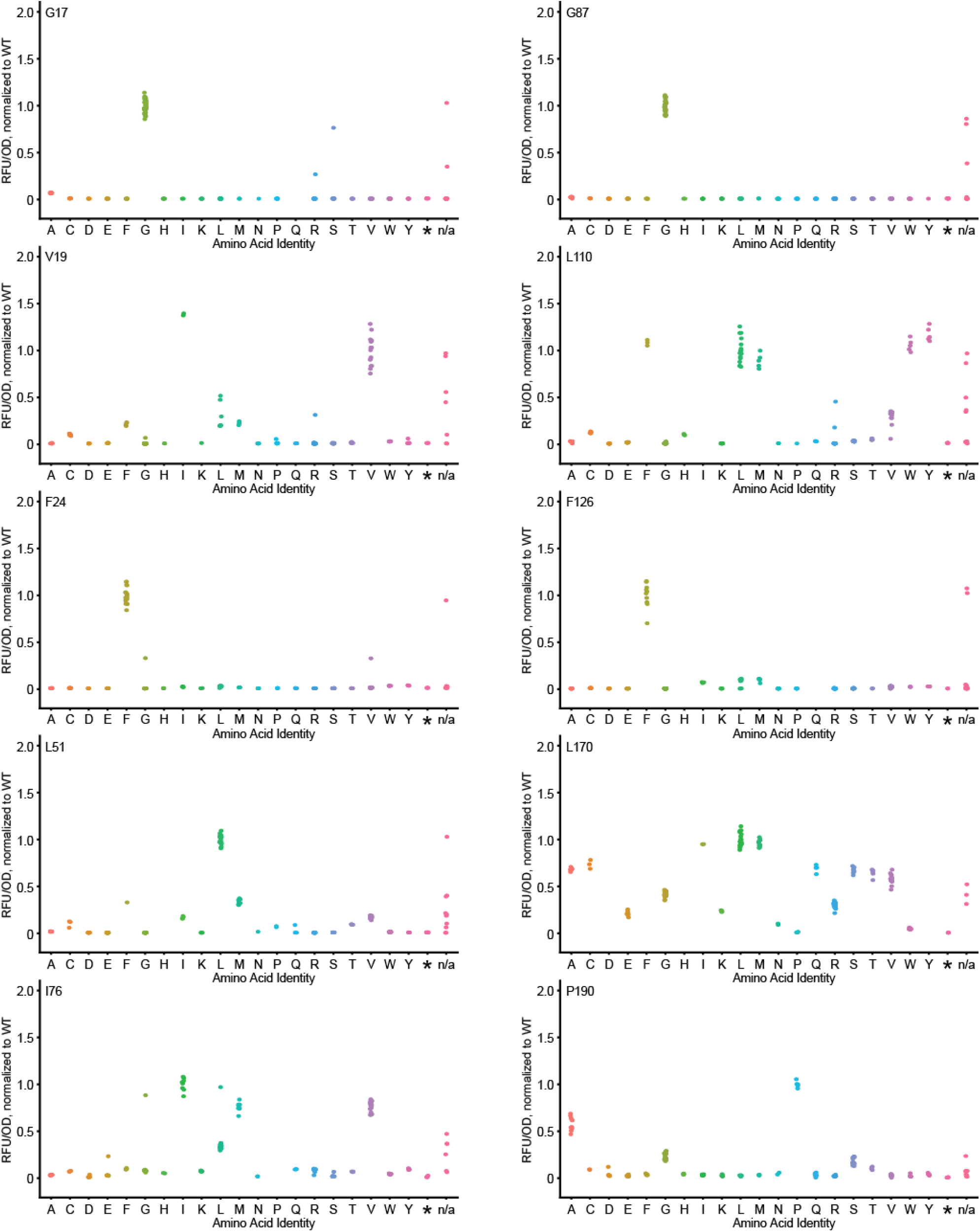
Fluorescence data for site-saturation libraries at favored residues in secBFP2.1 identified by the model. Raw fluorescence values were normalized to OD600 and to the average wild-type value. Outliers were identified through the OPTICS algorithm and removed. n/a represents variant calls that failed to meet the specified thresholds (see Methods).

**SI Figure 7:**
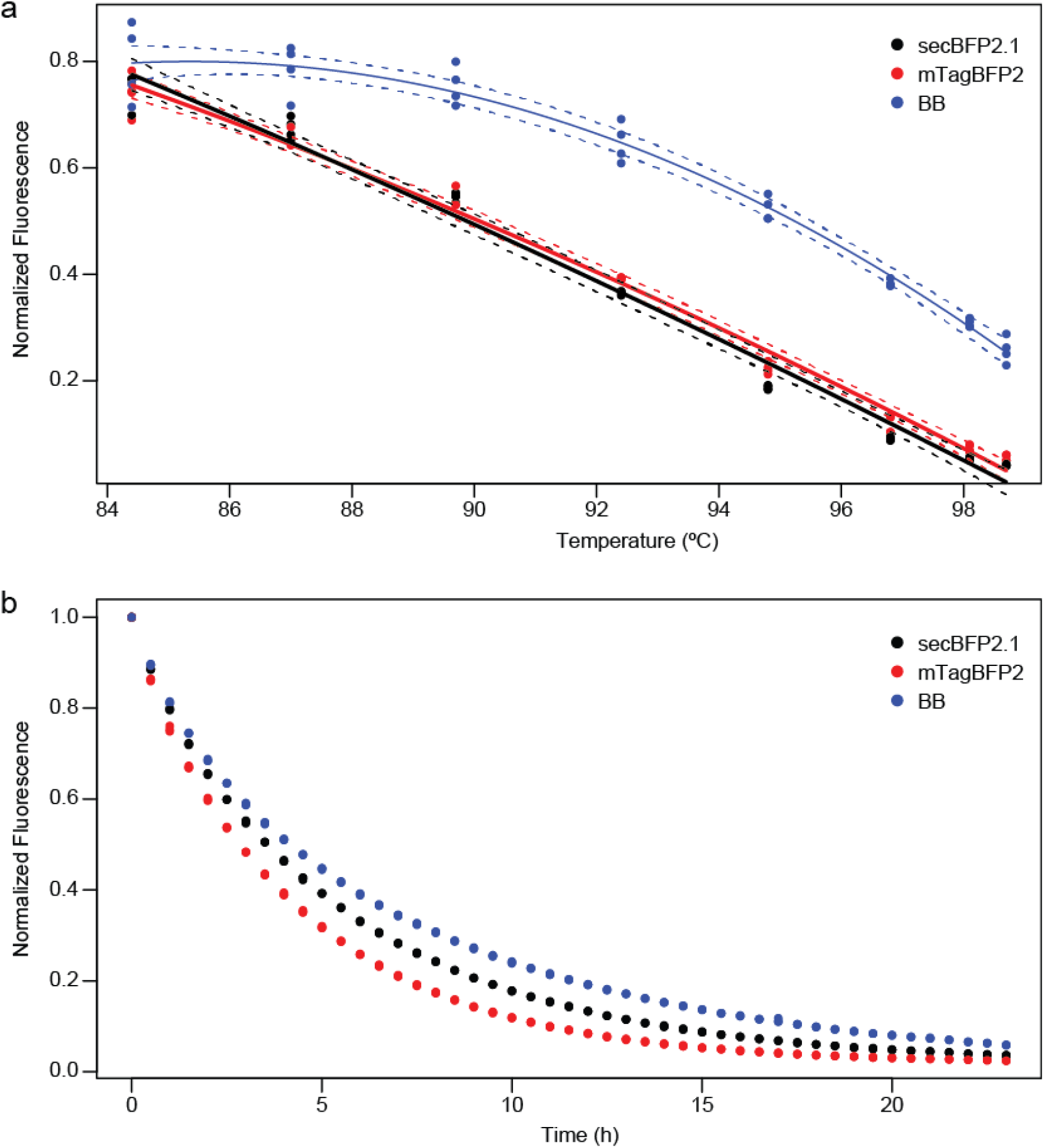
BFP-Bluebonnet (BB) exhibited improved folding compared to parental proteins. a) Plot of residual fluorescence after a ten minute thermal challenge at the indicated temperatures. Quadratic terms were significant in the global linear model and lines correspond to a quadratic model fit for each blue fluorescent protein. Dotted lines delineate 95% confidence intervals for the regression. mTagBFP2 and secBFP2.1 were not significantly different from each other while first order and quadratic terms for BB were significantly different compared to either parental protein. The assay was performed with 4-fold replication. b) Guanidinium melt of BFP variants. This assay was performed with 3-fold replication.

**Figure 8:**
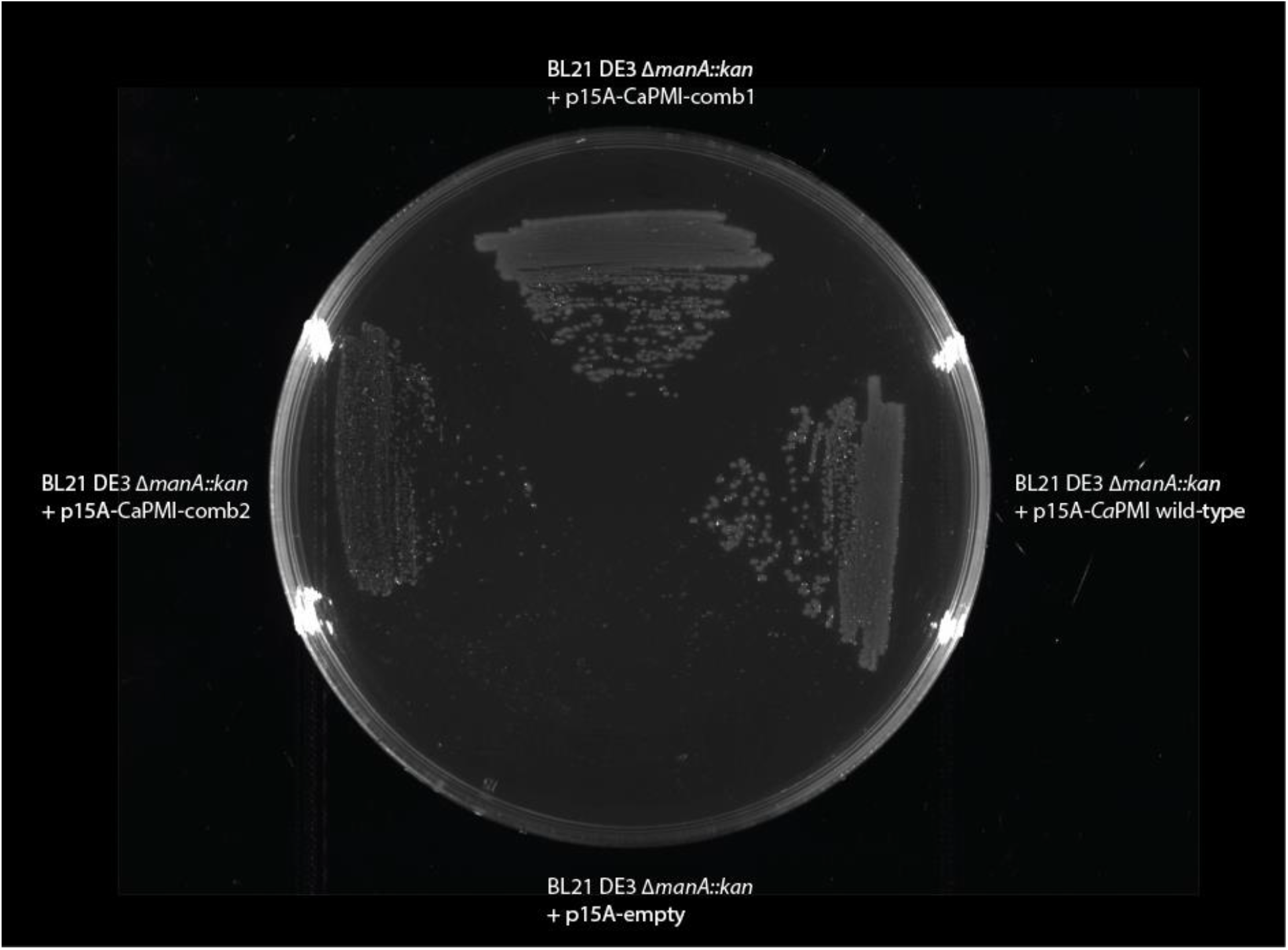
Mutant *Ca*PMI variants can complement deletion of the *E. coli manA* gene. *Ca*PMI-comb1 contains mutations D229W, N272K, L335A, N388S and S425T. *Ca*PMI-comb2 contains mutations S56A, G119A, Q157I, Q193D, D229T, C295V, L335E, K347R, S368N, K402R and Q428T. Growth of *Ca*PMI-comb2 was poorer than wildtype *Ca*PMI and *Ca*PMI-comb1.

**SI Figure 9:**
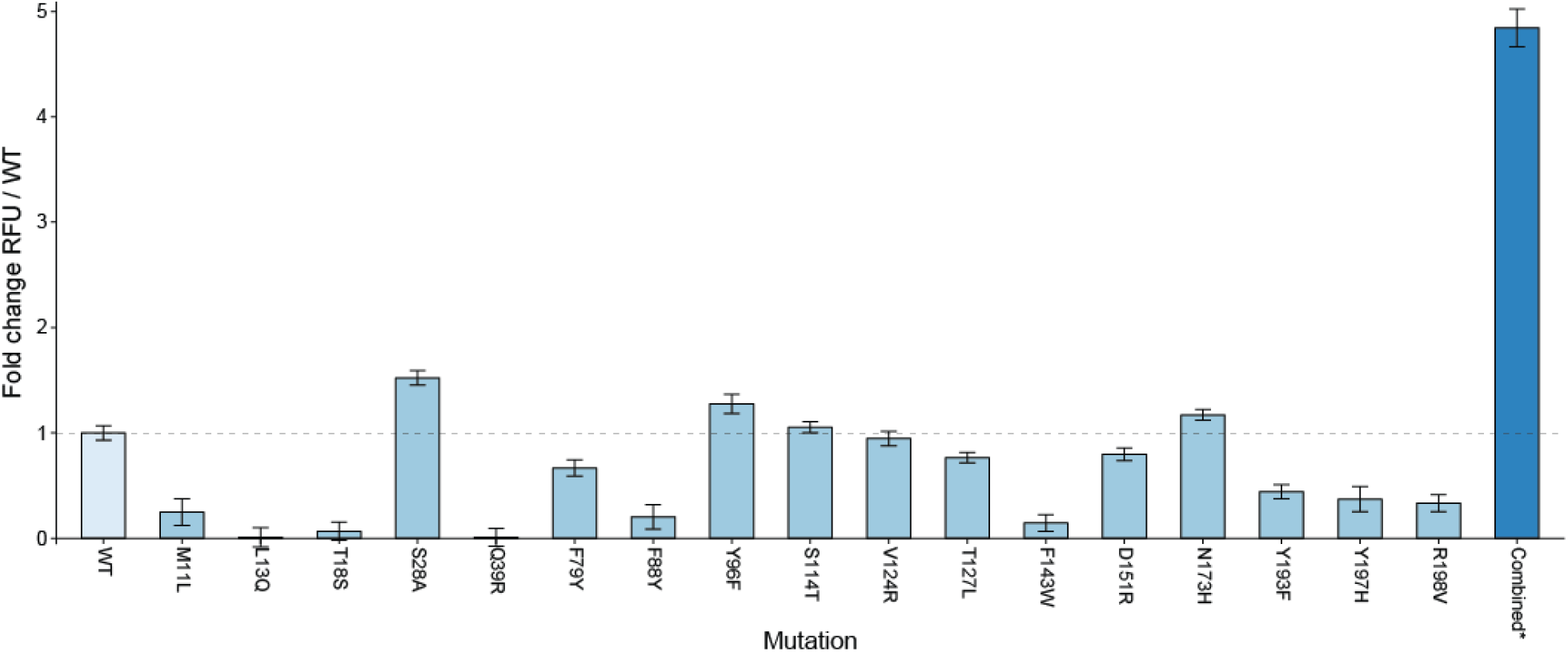
Fluorescence assay of secBFP2.1 variants containing a series of mutations predicted by the model. The combined BFP variant contains mutations S28A, S114T, T127L and N173H. While Y96F was identified as stabilizing by itself, it was deleterious in combination.

**Figure 10:**
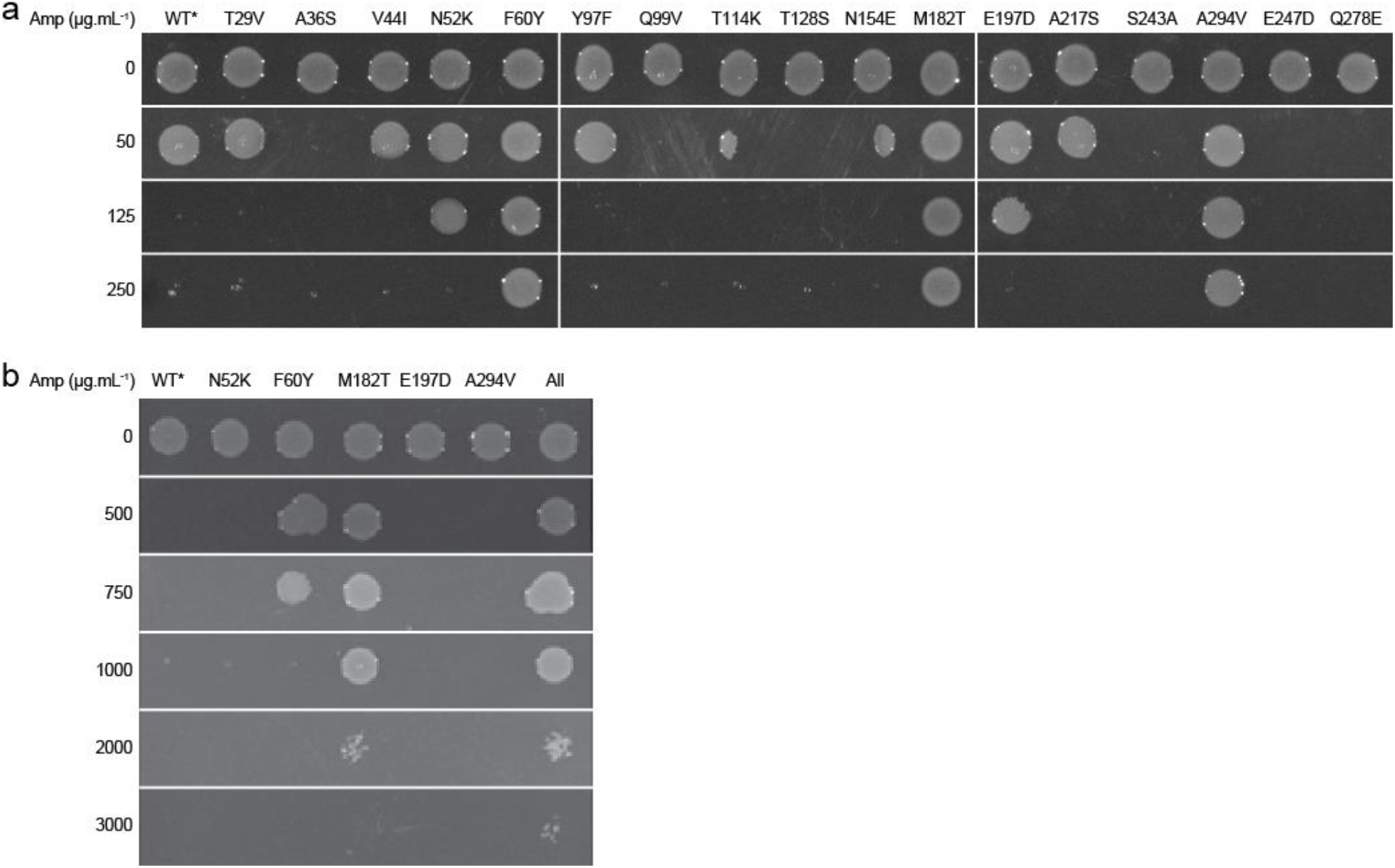
Antibiotic resistance assay of TEM-1 β-lactamase variants containing mutations predicted by the model. Individual mutants N52K, F60Y, M182T, E197D and A294V singularly and a combined variant containing all five stabilizing mutations resulted in increased ampicillin resistance. WT* contains the destabilizing mutation L250Q.

**Figure 11:**
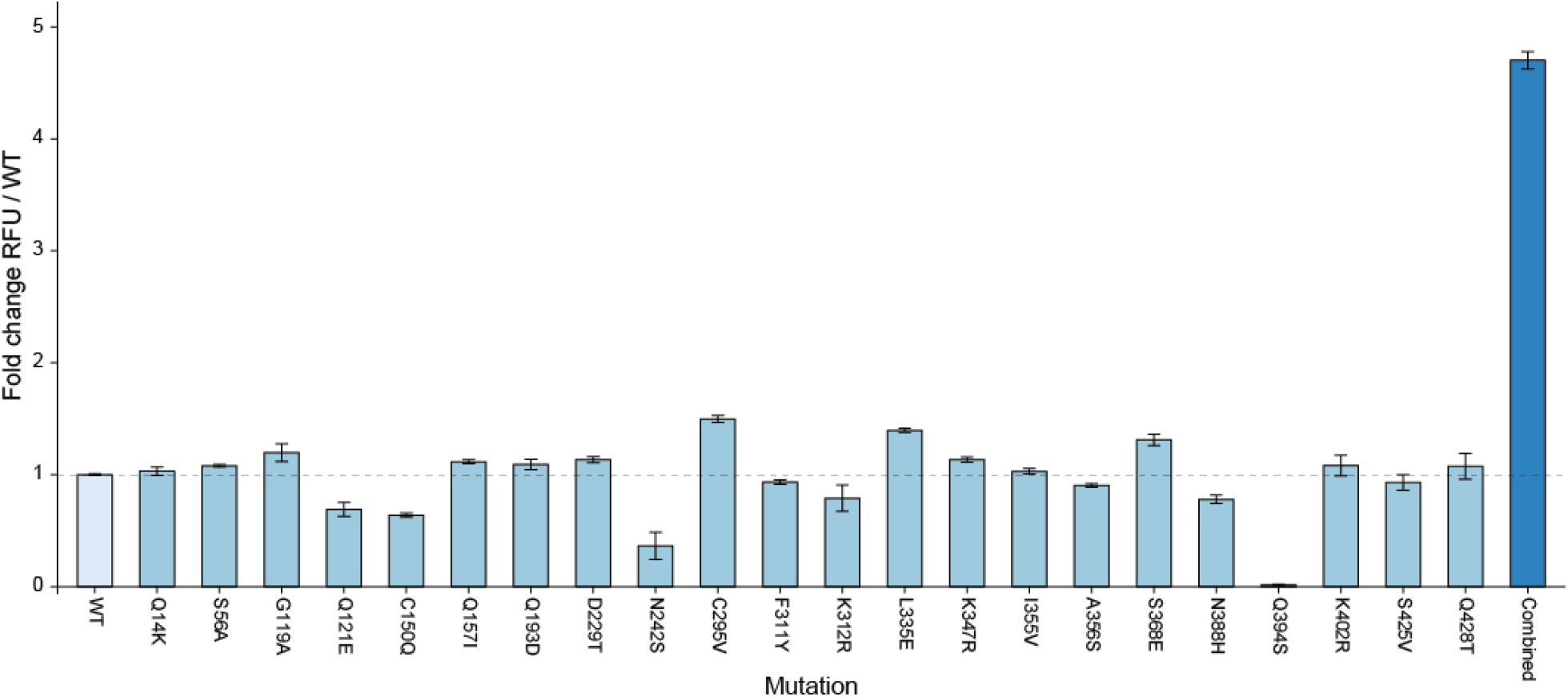
Fluorescence assay of the split-GFP-*Ca*PMI fusions containing a series of mutations predicted by the model. The combined *Ca*PMI variant contains mutations S56A, G119A, Q157I, Q193D, D229T, C295V, L335E, K347R, S368N, K402R and Q428T.

**SI Figure 12:**
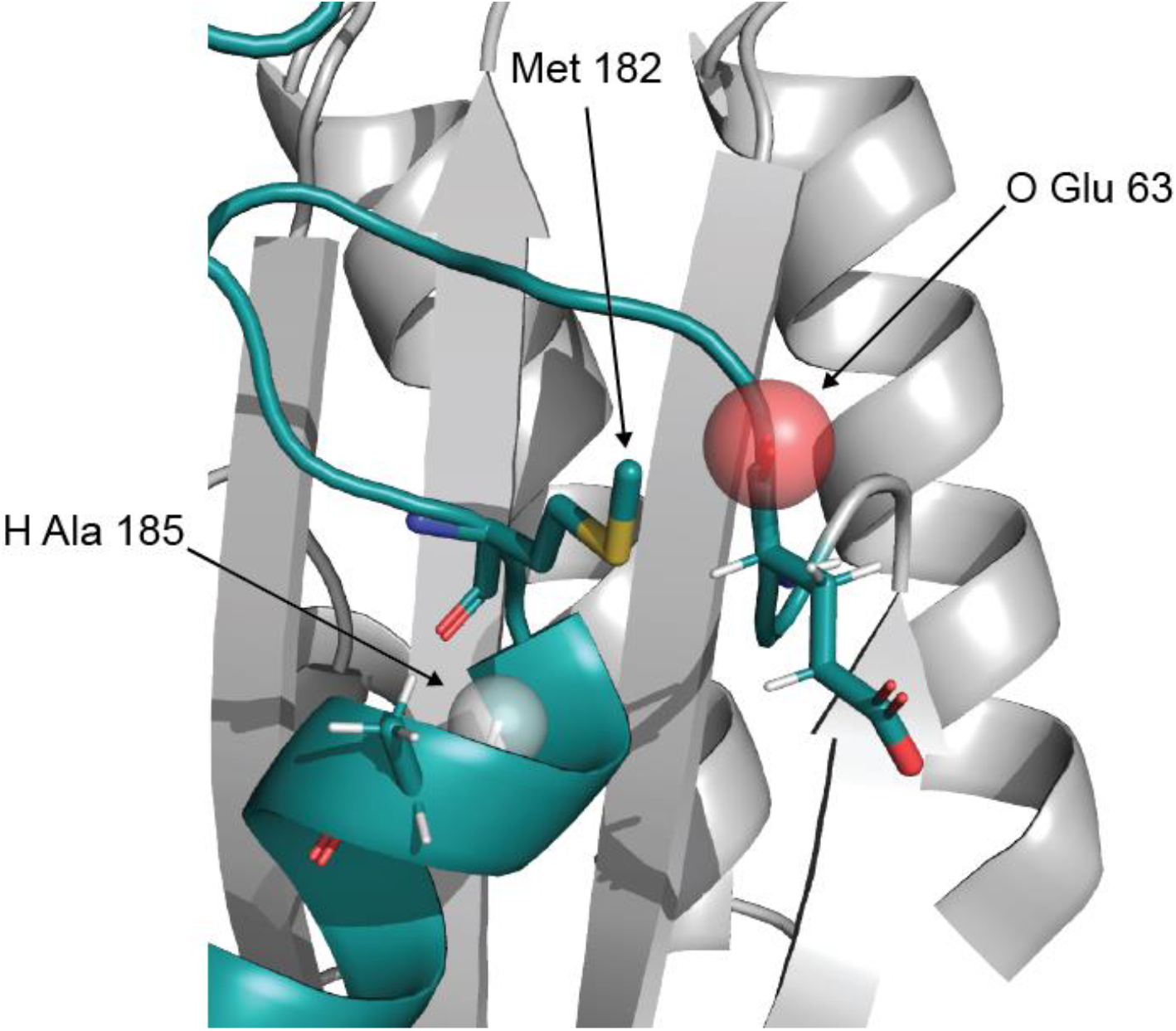
Masking of atoms reveals the mechanism of a global stabilizing mutation. Each atom in the Met 182 microenvironment was systematically deleted and the atoms favoring a mutation to threonine were identified. Of these, the two atoms, O Glu 63 and H Ala 185, change in probability by over 200-fold and have been identified previously in the literature as stabilization pathways for M182T^2^.

**SI Figure 13:**
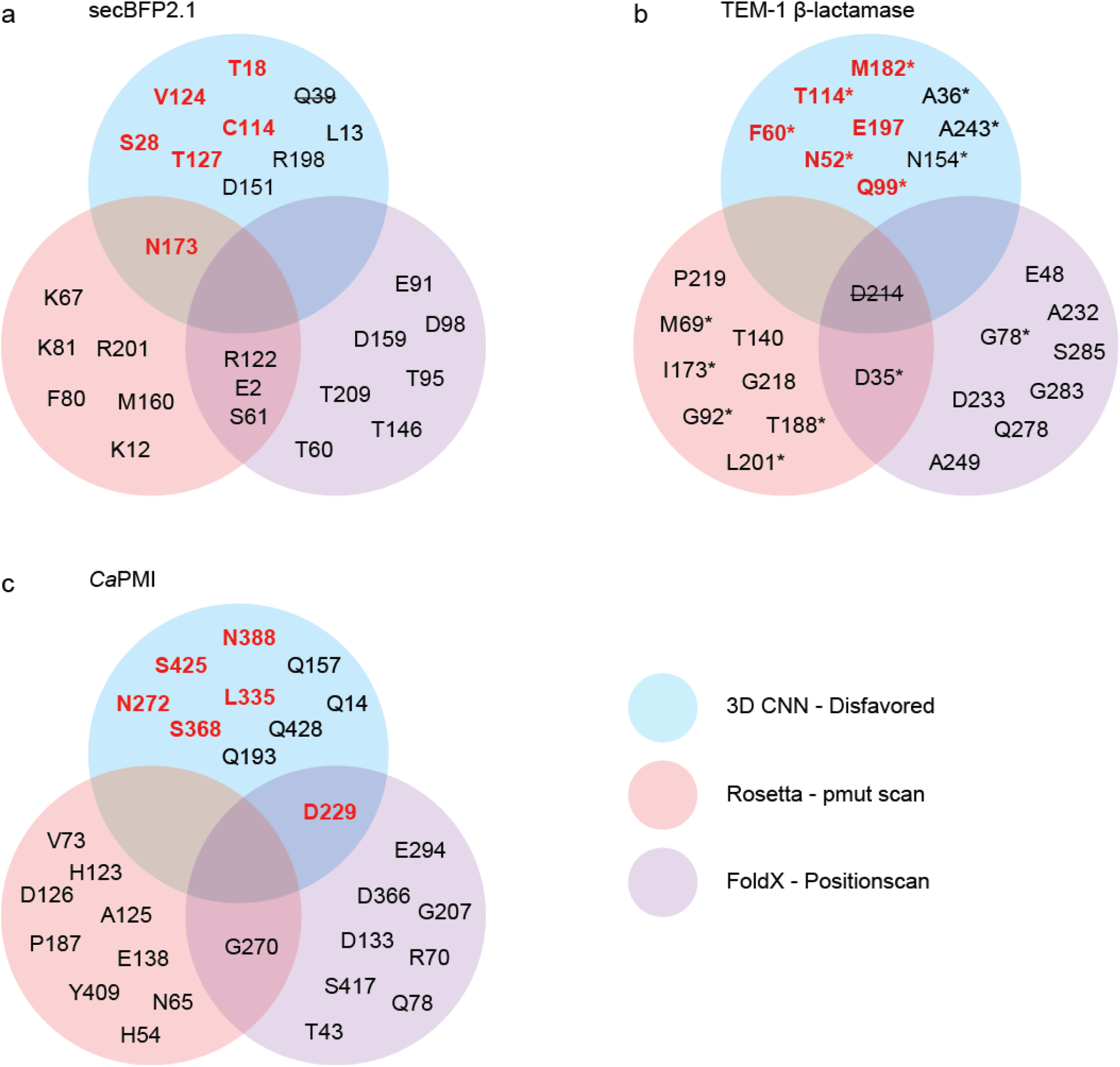
Venn diagram showing overlap between predictions of the 10 least favorable residues in model proteins made by different computational tools for protein design. Residues colored red indicate positions where we identified beneficial substitutions. Q39 in secBFP2.1 was not analyzed due repeated failure of our site-saturation library to assemble. D214 in TEM-1 β-lactamase was excluded due to its location in the enzyme active site. * Locations in TEM-1 β-lactamase where global suppressors and other beneficial substitutions have been identified^6, 7^.

### Supplementary Tables

**SI Table 1.**
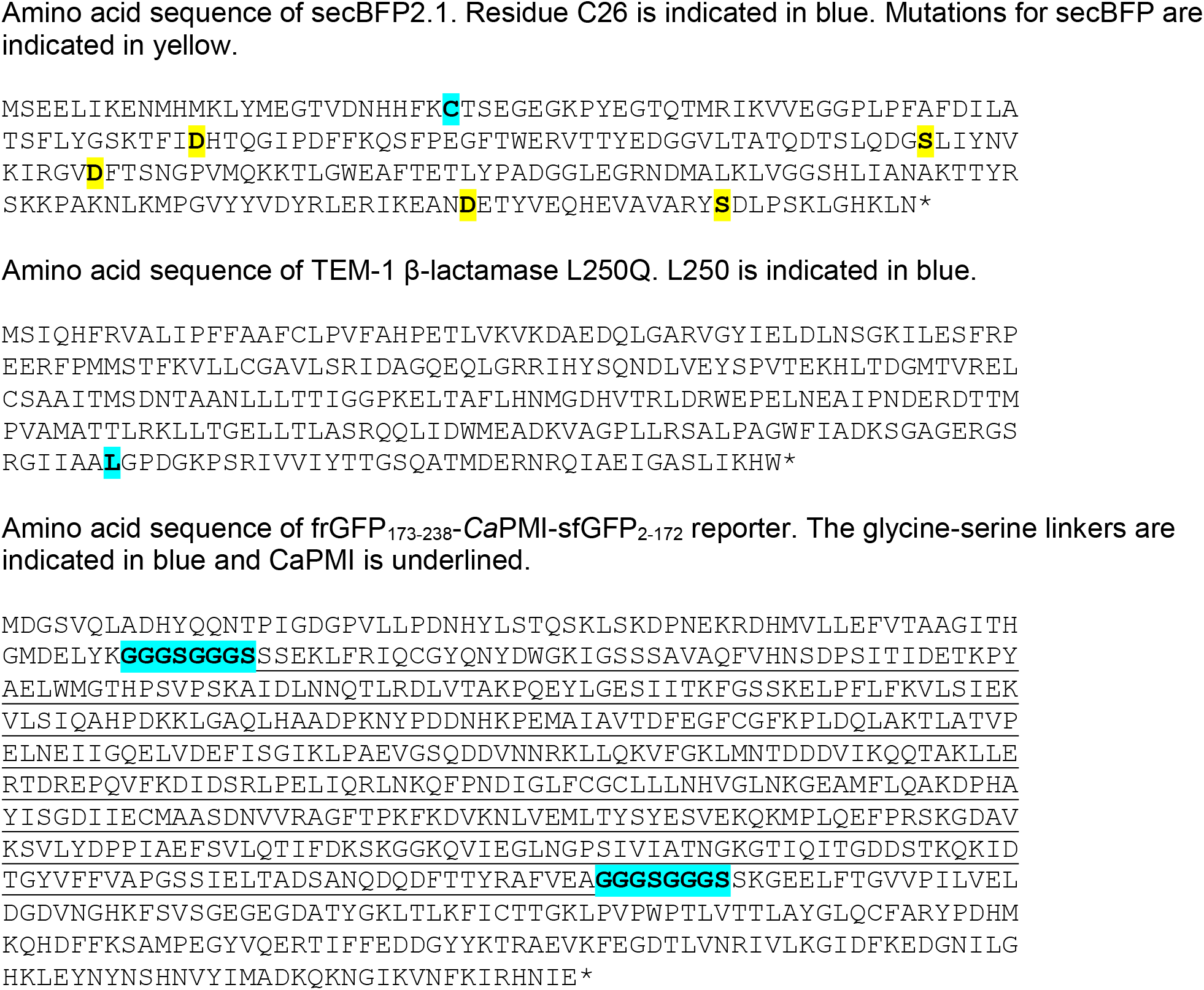
Amino acid sequences of model proteins.

## References

1. Jensen, P.F. et al. Structure and Dynamics of a Promiscuous Xanthan Lyase from Paenibacillus nanensis and the Design of Variants with Increased Stability and Activity. Cell chemical biology 26, 191–202 e196 (2019).

2. Windle, C.L. et al. Extending enzyme molecular recognition with an expanded amino acid alphabet. Proceedings of the National Academy of Sciences of the United States of America 114, 2610–2615 (2017).

3. Studer, S. et al. Evolution of a highly active and enantiospecific metalloenzyme from short peptides. Science (New York, N.Y.) 362, 1285–1288 (2018).

4. Trudeau, D.L., Kaltenbach, M. & Tawfik, D.S. On the Potential Origins of the High Stability of Reconstructed Ancestral Proteins. Molecular biology and evolution 33, 2633–2641 (2016).

5. Potapov, V., Cohen, M. & Schreiber, G. Assessing computational methods for predicting protein stability upon mutation: good on average but not in the details. Protein engineering, design & selection: PEDS 22, 553–560 (2009).

6. Jia, L., Yarlagadda, R. & Reed, C.C. Structure Based Thermostability Prediction Models for Protein Single Point Mutations with Machine Learning Tools. PloS one 10, e0138022 (2015).

7. Wu, Z., Kan, S.B.J., Lewis, R.D., Wittmann, B.J. & Arnold, F.H. Machine learning-assisted directed protein evolution with combinatorial libraries. Proceedings of the National Academy of Sciences of the United States of America 116, 8852–8858 (2019).

8. Alley, E.C., Khimulya, G., Biswas, S., AlQuraishi, M. & Church, G.M. Unified rational protein engineering with sequence-based deep representation learning. Nature methods (2019).

9. Torng, W. & Altman, R.B. 3D deep convolutional neural networks for amino acid environment similarity analysis. BMC bioinformatics 18, 302 (2017).

10. Joosten, R.P. et al. PDB_REDO: automated re-refinement of X-ray structure models in the PDB. Journal of applied crystallography 42, 376–384 (2009).

11. Gray, V.E., Hause, R.J., Luebeck, J., Shendure, J. & Fowler, D.M. Quantitative Missense Variant Effect Prediction Using Large-Scale Mutagenesis Data. Cell systems 6, 116–124 e113 (2018).

12. Wang, J., Cao, H., Zhang, J.Z.H. & Qi, Y. Computational Protein Design with Deep Learning Neural Networks. Sci Rep 8, 6349 (2018).

13. Li, Z., Yang, Y., Faraggi, E., Zhan, J. & Zhou, Y. Direct prediction of profiles of sequences compatible with a protein structure by neural networks with fragment-based local and energy-based nonlocal profiles. Proteins 82, 2565–2573 (2014).

14. O’Connell, J. et al. SPIN2: Predicting sequence profiles from protein structures using deep neural networks. Proteins 86, 629–633 (2018).

15. Costantini, L.M., Subach, O.M., Jaureguiberry-bravo, M., Verkhusha, V.V. & Snapp, E.L. Cysteineless non-glycosylated monomeric blue fluorescent protein, secBFP2, for studies in the eukaryotic secretory pathway. Biochemical and biophysical research communications 430, 1114–1119 (2013).

16. Zimmerman, M.I. et al. Prediction of New Stabilizing Mutations Based on Mechanistic Insights from Markov State Models. ACS central science 3, 1311–1321 (2017).

17. Bratulic, S., Gerber, F. & Wagner, A. Mistranslation drives the evolution of robustness in TEM-1 beta-lactamase. Proceedings of the National Academy of Sciences of the United States of America 112, 12758–12763 (2015).

18. Guthrie, V.B., Allen, J., Camps, M. & Karchin, R. Network models of TEM beta-lactamase mutations coevolving under antibiotic selection show modular structure and anticipate evolutionary trajectories. PLoS computational biology 7, e1002184 (2011).

## References

1. Joosten, R.P. et al. PDB_REDO: automated re-refinement of X-ray structure models in the PDB. Journal of applied crystallography 42, 376–384 (2009).

2. Dolinsky, T.J. et al. PDB2PQR: expanding and upgrading automated preparation of biomolecular structures for molecular simulations. Nucleic acids research 35, W522–525 (2007).

3. Mitternacht, S. FreeSASA: An open source C library for solvent accessible surface area calculations. F1000Research 5, 189 (2016).

4. Torng, W. & Altman, R.B. 3D deep convolutional neural networks for amino acid environment similarity analysis. BMC bioinformatics 18, 302 (2017).

5. Gray, V.E., Hause, R.J., Luebeck, J., Shendure, J. & Fowler, D.M. Quantitative Missense Variant Effect Prediction Using Large-Scale Mutagenesis Data. Cell systems 6, 116–124 e113 (2018).

6. Jacquier, H. et al. Capturing the mutational landscape of the beta-lactamase TEM-1. Proceedings of the National Academy of Sciences of the United States of America 110, 13067–13072 (2013).

7. Cabantous, S., Rogers, Y., Terwilliger, T.C. & Waldo, G.S. New molecular reporters for rapid protein folding assays. PloS one 3, e2387 (2008).

## Supplementary References

1. Kather, I., Jakob, R.P., Dobbek, H. & Schmid, F.X. Increased folding stability of TEM-1 beta-lactamase by in vitro selection. Journal of molecular biology 383, 238–251 (2008).

2. Zimmerman, M.I. et al. Prediction of New Stabilizing Mutations Based on Mechanistic Insights from Markov State Models. ACS central science 3, 1311–1321 (2017).

3. Farzaneh, S. et al. Implication of Ile-69 and Thr-182 residues in kinetic characteristics of IRT-3 (TEM-32) beta-lactamase. Antimicrobial agents and chemotherapy 40, 2434–2436 (1996).

4. Orencia, M.C., Yoon, J.S., Ness, J.E., Stemmer, W.P. & Stevens, R.C. Predicting the emergence of antibiotic resistance by directed evolution and structural analysis. Nature structural biology 8, 238–242 (2001).

5. Torng, W. & Altman, R.B. 3D deep convolutional neural networks for amino acid environment similarity analysis. BMC bioinformatics 18, 302 (2017).

6. Bratulic, S., Gerber, F. & Wagner, A. Mistranslation drives the evolution of robustness in TEM-1 beta-lactamase. Proceedings of the National Academy of Sciences of the United States of America 112, 12758–12763 (2015).

7. Guthrie, V.B., Allen, J., Camps, M. & Karchin, R. Network models of TEM beta-lactamase mutations coevolving under antibiotic selection show modular structure and anticipate evolutionary trajectories. PLoS computational biology 7, e1002184 (2011).

